# Seafloor Incubation Experiments at Deep-Sea Hydrothermal Vents Reveal Distinct Biogeographic Signatures of Autotrophic Communities

**DOI:** 10.1101/2023.07.20.549880

**Authors:** Heather Fullerton, Lindsey Smith, Alejandra Enriquez, David Butterfield, C. Geoffrey Wheat, Craig L. Moyer

**Affiliations:** Department of Biology, College of Charleston, Charleston, SC, United States; Department of Biology, Western Washington University, Bellingham, WA, United States; Cooperative Institute for Climate, Ocean, and Ecosystem Studies, University of Washington and NOAA/PMEL, Seattle, Washington, USA; Institute of Marine Studies, College of Fisheries and Ocean Sciences, University of Alaska Fairbanks, Moss Landing, CA, USA

**Keywords:** Kama’ehuakanaloa Seamount, hydrothermal vents, community structure, Zetaproteobacteria, Campylobacterota

## Abstract

The discharge of hydrothermal vents on the seafloor provides energy sources for dynamic and productive ecosystems, which are supported by chemosynthetic microbial populations. These populations use the energy gained by oxidizing the reduced chemicals contained within the vent fluids to fix carbon dioxide and support multiple trophic levels. Hydrothermal discharge is ephemeral and chemical composition of such fluids varies over space and time, which can result in geographically distinct microbial communities. To investigate the foundational members of the community, microbial growth chambers were placed within the hydrothermal discharge at Axial Seamount (Juan de Fuca Ridge), Magic Mountain Seamount (Explorer Ridge), and Kama’ehuakanaloa Seamount (Hawai’i hotspot). Campylobacteria were identified within the nascent communities, but different amplicon sequence variants were present at Axial and Kama’ehuakanaloa Seamounts, indicating that geography in addition to the composition of the vent effluent influences microbial community development. These results provide insights to nascent microbial community structure and shed light on the development of diverse lithotrophic communities at hydrothermal vents.

## Introduction

Deep-sea hydrothermal vents are dynamic and extremely productive biological ecosystems supported by chemosynthetic microbial populations (Jannasch and Mottl 1985). These ecosystems offer multiple unique circumstances for microbial growth, including substrate surrounding and near vent discharge, within the subsurface, plume emissions, and through the formation of symbiotic relationships with vent invertebrates (Karl, Wirsen and Jannasch 1980; Jannasch 1983; Murdock *et al*. 2021). The hydrothermal vents themselves provide energy to the microorganisms via the oxidation of reduced solutes that discharge from these habitats (Dick 2019). Overall, the metabolic capacity of the system depends on the chemical composition of the hydrothermal vent effluent (Karl 1995; Nakagawa and Takai 2008). The concentrations of reduced solutes can vary over space and time, and rapid changes in vent chemistry and temperature impact microbial community composition, structure, and function (Butterfield *et al*. 1997; Danovaro *et al*. 2017). Microbial species are often found at hydrothermal vents that are geographically separated; therefore, they are ideal systems for addressing questions of microbial biogeography and speciation (Price, Fullerton and Moyer 2015; Duchinski *et al*. 2019). At these habitats, there is evidence of dispersal as well as allopatric speciation in both bacteria and archaea (Price, Fullerton and Moyer 2015; Mino *et al*. 2017).

Bacteria within the Campylobacterota and Proteobacteria phyla are often found near hydrothermal vents. The Campylobacteria and Zetaproteobacteria are two of the major phylogenetic classes of Bacteria distributed below the subsurface, within the vent effluent, and near the vent orifice. Such bacteria form microbial biofilms, also referred to as microbial mats (Waite *et al*. 2017, 2018; Parks *et al*. 2018). Many of the dominant organisms at hydrogen- and sulfur-rich hydrothermal vents are grouped within the phylum Campylobacterota and class Campylobacteria (formerly Epsilonproteobacteria) with *Sulfurimonas, Sulfurovum, Nitritruptor, Thioreductor* and *Arcobacter* as the most common genera (Orcutt *et al*. 2011b). In contrast, at hydrothermal vent systems with high concentrations of reduced iron, the class Zetaproteobacteria is the dominant community member (Hager *et al*. 2017; Duchinski *et al*. 2019). They are the newest described class of Proteobacteria with one described genus, *Mariprofundus* (Emerson *et al*. 2007; Makita *et al*. 2017). Since their original identification at hydrothermal vents, Zetaproteobacteria have been shown to reside in specific ecological niches dispersed world-wide (McAllister *et al*. 2019).

Campylobacterota are found in diverse environments, have diverse metabolic strategies (Campbell *et al*. 2006) and in many hydrothermal vent ecosystems, members of the phylum Campylobacterota have been classified as the primary producers (Han and Perner 2015). Within this phylum, *Sulfurimonas* and *Sulfurovum* have been identified in active and inactive hydrothermal sulfides at the East Pacific Rise (Sylvan, Toner and Edwards 2012), microbial mats of the Mariana Arc and back-arc (Hager *et al*. 2017), vent effluent of the Mariana Arc and Axial Seamount (Huber *et al*. 2010; Akerman, Butterfield and Huber 2013) and hydrothermal sediments of the Mid-Atlantic Ridge (Cerqueira *et al*. 2018). *Sulfurimonas* spp. are able to grow with reduced sulfur compounds and hydrogen as their electron donor and aerobically with oxygen or anaerobically with nitrate or nitrite as their electron acceptors (Vetriani *et al*. 2014; Han and Perner 2015). *Sulfurimonas autotrophica* was isolated from the Mid-Okinawa Trough hydrothermal field and grows autotrophically with sulfide, elemental sulfur, and thiosulfate as electron donors (Inagaki *et al*. 2003). Genomic analysis of the globally-distributed *Sulfurimonas* shows evidence of allopatric speciation and suggests that geographic distance is the primary driver of genetic variation (Mino *et al*. 2017).

At iron-dominated venting locations, Zetaproteobacteria are considered as microbial ecosystem engineers (Hager *et al*. 2017). This means they have the potential to modulate the flow of organic carbon to other microbes and have the capacity to shape their habitat by producing iron oxyhydroxide minerals and exopolysaccharides, increasing richness and diversity within the microbial landscape, affecting the health and stability of the environment (Chan *et al*. 2011, 2016; Jesser *et al*. 2015; Hager *et al*. 2017). Even at low abundances, Zetaproteobacteria can impact their local environment and can also be considered a keystone taxa, because they are necessary for the survival of other microbes and are drivers of ecosystem biodiversity (Beam *et al*. 2018a). There are currently 59 OTUs of Zetaproteobacteria (zOTUs), four of which are considered cosmopolitan and another twelve to fifteen of which have high endemic abundances. The remaining zOTUs have rarely been observed (McAllister *et al*. 2019). A few isolates have been grown in pure culture including *Mariprofundus ferrooxydans* (strains PV-1 and JV-1) from Kama’ehuakanaloa Seamount and isolates from the estuarine water column of Chesapeake Bay, among others (Chiu *et al*. 2017). However, the cosmopolitan zOTUs, which are ubiquitous at iron-driven vent communities throughout the Pacific and Atlantic Oceans, have not yet been cultured (McAllister *et al*. 2011; Scott *et al*. 2015). Therefore, the physiology of these cosmopolitan zOTUs has been inferred from the distantly related isolates and metagenomics (Field *et al*. 2014; Fullerton *et al*. 2017; He *et al*. 2017).

A previous study using microbial growth chambers at Kama’ehuakanaloa Seamount (formerly known as Lo’ihi Seamount) identified Zetaproteobacteria as the dominant colonizers (Rassa *et al*. 2009). To date, microbiological studies at Axial and Magic Mountain have only been conducted on fluid samples to assess microbial vent-associated community structure (Meyer *et al*. 2013a; Anderson *et al*. 2017; Moulana *et al*. 2020), with the exception of an earlier pilot-study using T-RFLP analysis (Engebretson 2002). The microbial growth chambers (MGCs) provide an inert surface to examine nascent communities of lithotrophs to better understand how microbial communities rebuild and interact. To further investigate how microbial communities develop across geographically diverse hydrothermal vents, microbial growth chambers were deployed within the hydrothermal discharge at Axial Seamount (Juan de Fuca Ridge), Magic Mountain Seamount (Explorer Ridge), and Kama’ehuakanaloa Seamount (Hawai’i hotspot). The analysis of the nascent communities and comparisons across distinct hydrothermal vents addresses how microbial communities develop and how diversity is established within these complex ecosystems.

## Materials and Methods

### Construction of microbial growth chambers

Microbial growth chambers (MGCs) were constructed with three 3” sections of plexiglass cylinders that were enclosed on top and bottom with 202µm Nytex mesh to restrict grazing by macrofauna (Rassa *et al*. 2009). Each of the three chambers contained a total of ∼300g of hand-woven 8µm silica wool (yielding approximately 33 m^2^ of surface area) as a fresh surface to facilitate microbial attachment and growth. Negative buoyancy was achieved by addition of a stainless-steel eyebolt in the center of the three chambers, which also served as an attachment point for polyurethane line for ease of deployment and collection.

### Sample Collection

The MGCs were incubated from two to 19 days at venting locations at three different hydrothermal vent sites (1) Axial Seamount, (2) Magic Mountain and (3) Kama’ehuakanaloa Seamounts. Vent locations and deployment and *in situ* incubation times are summarized in Table 1. At Kama’ehuakanaloa, MGCs were deployed using the submersible *PISCES V* and the remotely operated vehicle (ROV) *JASON* in 1998, 2004 and 2009. At Magic Mountain, MGCs were deployed and collected with the ROV *ROPOS* in 2002. At Axial, MGCs were deployed using the ROV *ROPOS* in 1998, 1999, 2000 and 2002. In all cases, temperatures were monitored by either HOBO miniature temperature recorders placed at the same location as the MGC or by a temperature probe attached to the manipulator of the ROV at the time of MGC deployment and recovery. Upon recovery, each MGC was placed in a sealed box at the seafloor to minimize disturbance and flushing of the chambers during assent and recovery of the vehicle. Immediately upon recovery, MGCs were aseptically penetrated and silica wool removed, placed into heavyweight Ziploc-style sample bags, and immediately frozen. The frozen samples were shipped and stored at −80°C until further processing in the laboratory.

**Table 1.** *In situ* incubation times, location, cruise year, marker, and temperature of incubation for each microbial growth chamber (MGC).

| MGC | Duration<br>(days) | Location | Cruise<br>Year | Marker | Temperature<br>(°C) | Observed | Simpson |
| --- | --- | --- | --- | --- | --- | --- | --- |
| AxMGC-01 | 2 | Axial Caldera | 1998 | N4 | 23.1 | 117 | 0.66 |
| AxMGC-03 | 4 | Axial Caldera | 1998 | 113 | 22.3 | 1278 | 0.95 |
| AxMGC-05 | 16 | Axial Caldera | 1998 | 33 | 28.2 | 102 | 0.91 |
| AxMGC-08 | 2 | Axial Caldera | 1998 | 33 | 28.2 | 840 | 0.81 |
| AxMGC-10 | 14 | Axial Caldera | 1998 | 33 | 28.2 | 186 | 0.95 |
| AxMGC-18 | 12 | Axial Caldera | 1998 | N2 | 8.0 | 861 | 0.86 |
| AxMGC-36 | 5 | Axial Caldera | 1999 | N4 | 17.0 | 983 | 0.71 |
| AxMGC-38 | 5 | Axial Caldera | 1999 | N4 | 17.0 | 626 | 0.68 |
| AxMGC-43 | 5 | Axial Caldera | 1999 | 33 | 63.8 | 449 | 0.91 |
| AxMGC-44 | 7 | Axial Caldera | 2000 | 33 | 26.0 | 617 | 0.95 |
| AxMGC-47 | 7 | Axial Caldera | 2000 | N4 | 15.0 | 843 | 0.96 |
| AxMGC-66 | 19 | Axial Caldera | 2002 | N6 | 8.5 | 1144 | 0.97 |
| ExMGC-03 | 4 | Magic Mountain | 2002 | 73 | 21.0 | 544 | 0.94 |
| ExMGC-04 | 4 | Magic Mountain | 2002 | 81 | 11.5 | 140 | 0.95 |
| LMGC-L7 | 4 | Kama‘ehuakanaloa | 1998 | 20 | 24.0 | 796 | 0.94 |
| LMGC-L9A | 4 | Kama‘ehuakanaloa | 1998 | 11 | 77.0 | 207 | 0.90 |
| LMGC-L10 | 4 | Kama‘ehuakanaloa | 1998 | 11 | 64.0 | 339 | 0.87 |
| LMGC-03 | 4 | Kama‘ehuakanaloa | 2004 | 29 | 18.0 | 1791 | 0.94 |
| LMGC-04 | 4 | Kama‘ehuakanaloa | 2004 | 30 | 60.0 | 1012 | 0.93 |
| LMGC-05 | 4 | Kama‘ehuakanaloa | 2004 | 38 | 44.0 | 947 | 0.92 |
| LMGC-07 | 4 | Kama‘ehuakanaloa | 2004 | 38 | 44.0 | 1534 | 0.93 |
| LMGC-08 | 4 | Kama‘ehuakanaloa | 2004 | 39 | 46.0 | 1022 | 0.94 |
| LMGC-09 | 4 | Kama‘ehuakanaloa | 2004 | 36 | 54.0 | 1728 | 0.95 |
| LMGC-10 | 4 | Kama‘ehuakanaloa | 2004 | 36 | 54.0 | 996 | 0.89 |
| LMGC-89 | 5 | Kama‘ehuakanaloa | 2009 | 39 | 47.5 | 202 | 0.84 |
| LMGC-90 | 8 | Kama‘ehuakanaloa | 2009 | 57 | 12.5 | 1382 | 0.82 |
| LMGC-91 | 5 | Kama‘ehuakanaloa | 2009 | 39 | 46.0 | 107 | 0.93 |
| LMGC-92 | 9 | Kama‘ehuakanaloa | 2009 | 57 | 24.3 | 1074 | 0.77 |
| LMGC-93 | 8 | Kama‘ehuakanaloa | 2009 | 36 | 35.6 | 242 | 0.95 |
| LMGC-94 | 8 | Kama‘ehuakanaloa | 2009 | 57 | 25.1 | 931 | 0.82 |
| LMGC-96 | 8 | Kama‘ehuakanaloa | 2009 | 38 | 50.2 | 129 | 0.94 |
| LMGC-97 | 9 | Kama‘ehuakanaloa | 2009 | 57 | 25.1 | 756 | 0.84 |
| LMGC-98 | 9 | Kama‘ehuakanaloa | 2009 | 34 | 50.6 | 281 | 0.96 |

For the Axial and Magic Mountain locations, hydrothermal discharge was collected with the hydrothermal fluid and particle sampler. An aliquot was analyzed on board for total hydrogen sulfide and pH using standard colorimetric and electrode techniques (Butterfield *et al*. 2004). An aliquot was acidified and analyzed ashore for dissolved iron and manganese by atomic absorption. Similarly, hydrothermal fluids that discharged from the Kama’ehuakanaloa Seamount were collected using Walden-Weiss titanium syringe (“Major”) samplers (Von Damm *et al*. 1985). Aliquots were measured at sea for total hydrogen sulfide and pH using the same methods as described previously (Butterfield *et al*. 2004). Dissolved Mn and Fe were measured ashore using standard inductively coupled plasma optical emission spectrometry (ICPOES) technique (Wheat *et al*. 2017).

### Isolation of microbial biomass

Silica wool was thawed on ice and then washed with sterile 1X PBS at 4°C for 30 min in sterile mason jars on a rotating platform. This wash solution was then decanted into 50 ml conical tubes for centrifugation at 6000xg for 15 min at 4°C. The supernatant was returned to the mason jar for another 1X PBS wash and the cell pellet was stored at 4°C. This process was completed until the 1X PBS was clear. Cell pellets were combined, then aliquoted into approximately 0.5 g wet weight for DNA extraction.

### DNA Extractions, Sequencing and Sequence Processing

Genomic DNA was extracted from cell pellets using the Fast DNA SPIN Kit for Soil (MP Biomedicals, Santa Ana, CA) according to the manufacturers’ protocol with a minor modification in which the gDNA was eluted in 1.0 mM Tris with 0.01 mM EDTA at pH 8. Cell lysis was optimized using two rounds of bead beating for 45 sec at a power setting of 5.5 using the FastPrep instrument (MP Biomedicals) with samples being placed on ice between runs. Extracted DNA was quantified by a Qubit 2.0 fluorometer using high-sensitivity reagents (ThermoFisher Scientific, Waltham, MA).

The V3-V4 regions of the SSU rRNA gene were amplified via polymerase chain reaction (PCR) from all MGC samples using bacterial primers 340F and 784R (Klindworth *et al*. 2013; Hager *et al*. 2017). The resulting amplicons were sequenced using a MiSeq (Illumina, San Diego, CA) as per manufacturer’s protocol generating 2 x 300bp paired-end reads. The resulting reads were trimmed of primers using CutAdapt (Martin 2011). The trimmed reads were then processed using the Divisive Amplicon Denoising Algorithm 2 (DADA2) v1.26.0 with pseudopooling following the previously described protocols (Lee *et al*. 2015; Callahan *et al*. 2016; Callahan, McMurdie and Holmes 2017) with R version 4.01 and using the Silva v138 database for assigning taxonomy. Samples were normalized using a center log-ratio transformation as implemented with microbiome version 1.10.0 in R (Lahti and Shetty 2017). Further analysis was completed using phyloseq version 1.32 (McMurdie and Holmes 2013). Analysis of Zeta OTUs was completed on SILVA-aligned Zetaproteobacteria identified sequences with ZetaHunter (McAllister, Moore and Chan 2018). The network was developed using the top 0.5% amplicon sequence variants (ASVs) by mean relative abundance, a total of 41 ASVs. Network construction and analysis were performed using the NetCoMi package for R (Peschel *et al*. 2021), employing the −propr” measure for estimating associations (Lovell *et al*. 2015; Erb and Notredame 2016; Erb *et al*. 2017; Quinn *et al*. 2017, 2018, 2019) and the −SPRING” method for visualizing associations (Yoon, Gaynanova and Müller 2019) and an eigenvector centrality cutoff of 0.95.

### Data Availability Statement

All sequence data are available through the NCBI Sequence Read Archive study number SUB11788075 (BioProject: PRJNA858068). Hydrothermal fluid compositions are provided in Table 2.

**Table 2.** Geochemical profiles associated with the MGCs.

| MGC | pH | H<sub>2</sub>S $\mu\text{mol/kg}$ | dFe $\mu\text{mol/kg}$ | Mn $\mu\text{mol/kg}$ |
| --- | --- | --- | --- | --- |
| AxMGC-01 | 5.55 | 0.76 | 76.50 | 2.50 |
| AxMGC-03 | 5.73 | 0.41 | 7.50 | 3.50 |
| AxMGC-05 | 5.04 | 1.79 | 1.58 | 17.75 |
| AxMGC-08 | 5.04 | 1.79 | 1.58 | 17.75 |
| AxMGC-10 | 5.04 | 1.79 | 1.58 | 17.75 |
| AxMGC-18 | 6.22 | 0.07 | 51.00 | 26.00 |
| AxMGC-36 | 5.99 | 2.35 | 0.31 | 0.01 |
| AxMGC-38 | 5.99 | 2.35 | 0.31 | 0.01 |
| AxMGC-43 | 4.86 | 2.12 | 1.86 | 29.88 |
| AxMGC-44 | 5.98 | 0.15 | 1.97 | 19.70 |
| AxMGC-47 | 5.95 | 0.08 | 15.20 | 7.70 |
| AxMGC-66 | 6.89 | 5.03 | 6.77 | 4.60 |
| ExMGC-03 | 5.48 | 315.48 | 11.80 | 104.30 |
| ExMGC-04 | 5.29 | 559.24 | 20.10 | 89.30 |
| LMGC-03 | 6.36 | 0.02 | 428.20 | 9.66 |
| LMGC-04 | 5.92 | 0.00 | 59.60 | 1.15 |
| LMGC-05 | 5.82 | 0.00 | 110.00 | 2.52 |
| LMGC-07 | 5.82 | 0.00 | 110.00 | 2.52 |
| LMGC-08 | 5.90 | 1.77 | 414.93 | 15.57 |
| LMGC-09 | 6.32 | 0.01 | 75.10 | 3.47 |
| LMGC-10 | 6.32 | 0.01 | 75.10 | 3.47 |
| LMGC-L10 | 5.37 | 3.85 | 46.75 | 40.76 |
| LMGC-L7 | 5.96 | 0.00 | 27.92 | 5.84 |
| LMGC-L9A | 5.37 | 3.85 | 46.75 | 40.76 |

## Results

### Site Descriptions

MGCs at the time of placement and recovery of representative short-term incubations are shown in Figure 1. At Kama’ehuakanaloa, Hiolo North is represented by Markers 31, 36 and 39, Hiolo South is represented by Markers 34 and 38 and Pohaku is represented by Marker 57. Both MGCs from Explorer Ridge were incubated at Magic Mountain. At Axial, Cloud Vent is represented by Markers N2 and N4. Location for these markers have been fully described (Glazer and Rouxel 2009; Opatkiewicz, Butterfield and Baross 2009; Deschamps *et al*. 2013; Fullerton *et al*. 2017).

**Figure 1.**
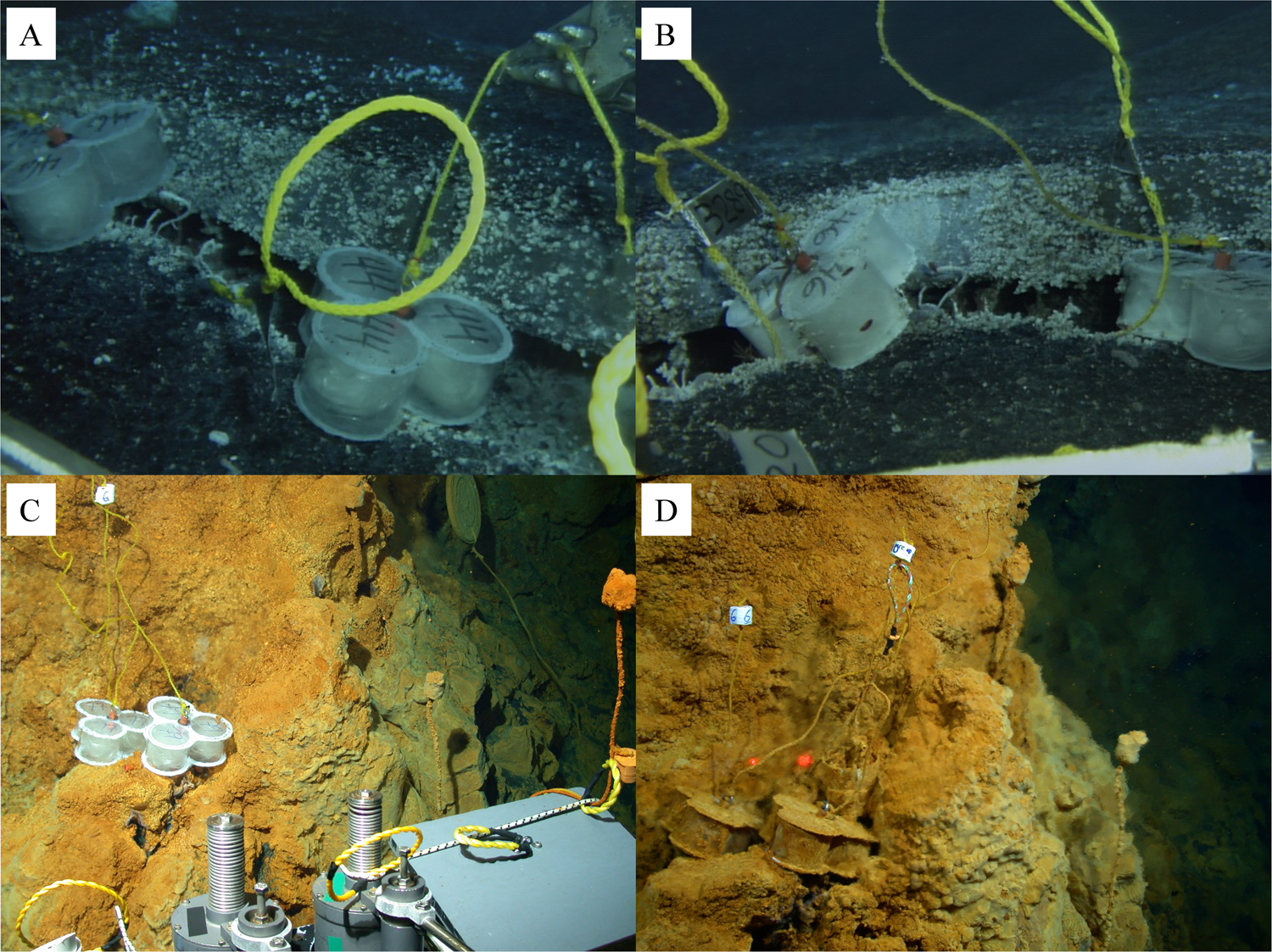
Pictures of representative MGCs before and after incubation showing development of microbial biomass as indicated in changes of opacity and color. (A) AxMGC.44 and AxMGC.46 at time of placement. (B) AxMGC.44 and AxMGC.46 at recovery after seven days. (C) LMGC.77 and LMGC.79 at time of placement. (D) LMGC.77 and LMGC.79 at time of recovery after 13 days.

In total, 33 MGCs were deployed and collected from the three locations, with two from Magic Mountain, twelve from Axial and 19 from Kama’ehuakanaloa (Table 1). Each MGC was placed within the fluid flow at each venting location and all vent sites had either an abundance of iron- or sulfur-dominated mats. During the *in situ* incubations, temperatures at Axial ranged from 8.0°C to 63.8°C and at Magic Mountain temperatures ranged from 11.5°C to 21°C. At Kama’ehuakanaloa, temperatures ranged from 12.5°C to 77°C (Table 1).

Although these sites have been relatively well sampled, the timeframes for chemistry sampling do not align with the MGC incubation timeframes for all the samples. In general, though, Kama’ehuakanaloa is characterized by lower temperature vents that have high concentrations of dissolved iron. Of the sampling locations at Kama’ehuakanaloa, Pohaku (Marker 57) has been routinely observed to have the highest abundance of Zetaproteobacteria. In 1996, an eruptive event at Kama’ehuakanaloa formed Pele’s Pit and high temperature venting fluids were observed shortly after. Therefore, the MGCs that were deployed in 1998 were incubated at the warmest of the observed temperatures at Kama’ehuakanaloa. Two sampling events shortly after the eruptive event recorded a decrease in thermal and fluid flux over eleven months (Wheat *et al*. 2000). Geochemistry sampling events in 2006, 2007 and 2008, showed Kama’ehuakanaloa had returned to a pre-eruption hydrothermal fluid composition <u>(Glazer and Rouxel 2009)</u>. However, during the 2009 collections at Kama’ehuakanaloa, no fluid compositions were determined.

For Axial and Magic Mountain MGC incubation periods, pH and concentrations of hydrogen sulfide, total dissolved iron, and manganese were determined, and temperatures were measured. At Kama’ehuakanaloa, these measurements were only made for the 1998 and 2004 incubation periods (Table 2). The pH at Axial ranged from 4.86 to 6.89, which overlaps in the ranges for both Magic Mountain and Kama’ehuakanaloa. The concentrations of hydrogen sulfide were lowest at Kama’ehuakanaloa, and highest at Magic Mountain. At Kama’ehuakanaloa, the MGCs that were incubated after the eruptive event had the highest concentration of hydrogen sulfide. Concentrations of dissolved manganese were overall highest at Magic Mountain and two of the post-eruption MGCs from Kama’ehuakanaloa. Dissolved iron was overall highest for the Kama’ehuakanaloa MGCs compared to the two highest concentrations measured in discharge where the Axial MGCs were deployed, even post-eruption.

### Sequencing and Community Structure

A total of 11,451,454 paired-end sequences were used as the input into DADA2 for quality filtering and removal of chimeras which resulted in a total of 7,611,025 contigs remaining representing 9,598 amplicon sequence variants (ASVs).

There was no pattern of diversity detected as related to each Seamount, and increasing duration did not correspond to increased diversity. For example, the observed ASVs were lowest in MGCs AxMGC-05, which was incubated for 16 days at Axial Seamount, and in LMGC-91, which was incubated for 5 days at Kama’ehuakanaloa Seamount. In these two MGCs there were 102 ASVs and 107, respectively (Table 1). The Chao1 and ACE species richness estimators show similar patterns to overall observed ASVs (Supplemental Table 1). Both alpha diversity estimators show LMGC-03, which was incubated Kama’ehuakanaloa for four days, to have the greatest count, with 1791 ASVs. The two MGCs from Magic Mountain, ExMGC-03 and ExMGC-04, were on the lower end with 544 and 140 observed ASVs. The MGCs with the lowest number of ASVs are not the most even as calculated by the Simpson’s index. MGC AxMGC-66 had the highest Simpson’s value, but also had 1158 ASVs by the Chao1 estimator. Conversely, MGC AxMGC-01 had the lowest Simpson’s value, and the third fewest ASVs by the Chao1 estimator.

Campylobacteria was the dominant class identified in MGCs incubated at Axial and Magic Mountain whereas Zetaproteobacteria was the dominant class at Kama’ehuakanaloa. As shown in Figure 2, the class Campylobacteria was identified throughout all Axial and Magic Mountain MGCs but only found in 11 of the 19 MGCs from Kama’ehuakanaloa. Alphaproteobacteria and Gammaproteobacteria were abundant at Kama’ehuakanaloa and largely absent within the Axial and Magic Mountain MGCs. Interestingly, the three Kama’ehuakanaloa MGCs, LMGC-L9A, LMGC-L7 and LMGC-L10 incubated shortly after the eruptive event show the lowest abundance of Zetaproteobacteria and highest abundance of Aquificae and Campylobacteria. Bacteroidia were distributed throughout MGCs from all three hydrothermal vent locations. The phylum Aquificota was present in only two MGCs from Axial, and one of the Magic Mountain MGCs. In the post-eruption MGCs incubated at Kama’ehuakanaloa, Aquificota were as abundant as the Campylobacteria (Figure 2).

**Figure 2.**
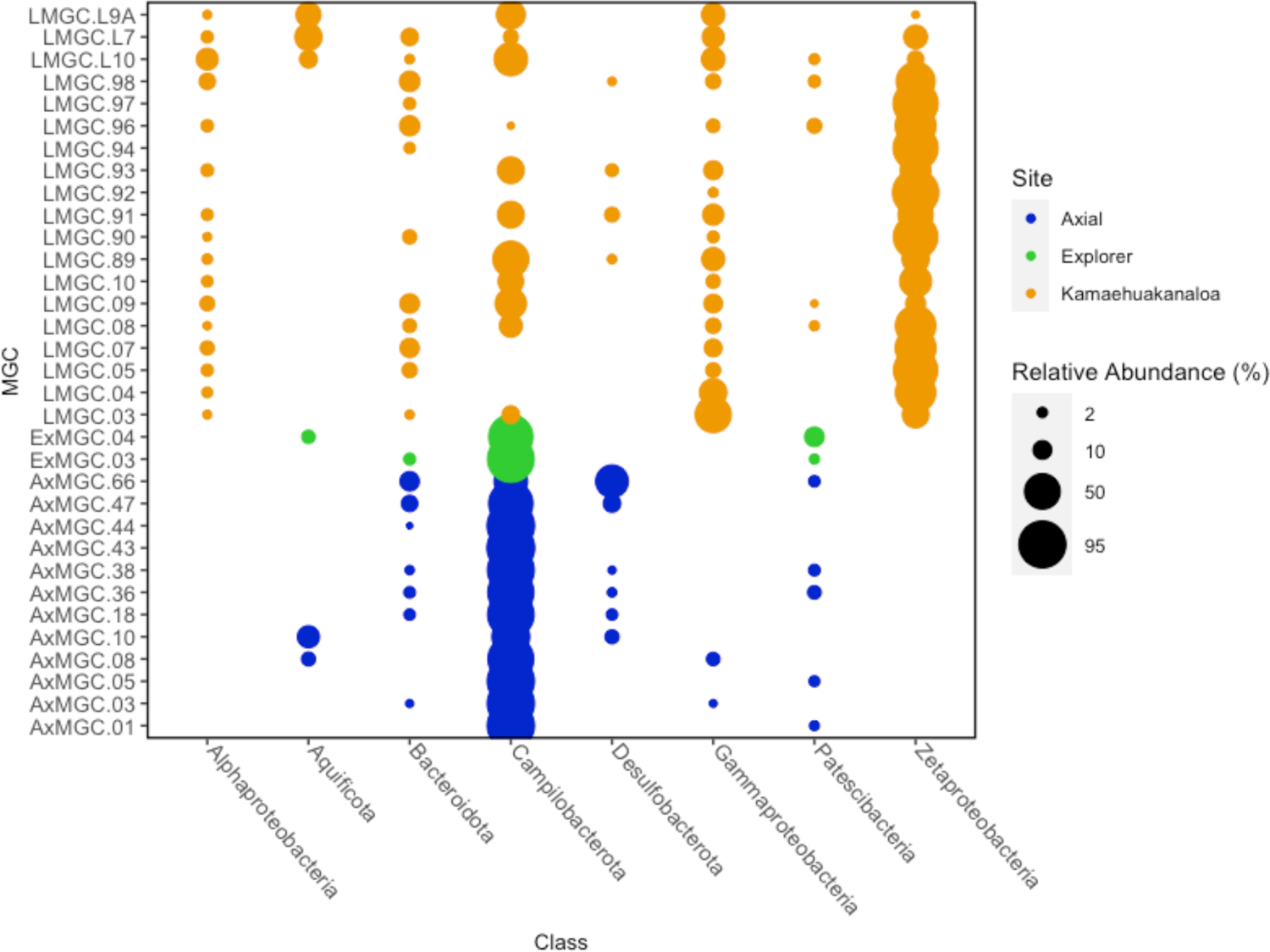
MGC community taxonomic distribution of associated bacteria genera of the top 1% in abundance.

The bacterial communities loosely grouped by marker location and seamount and did not group in the dendrogram by *in situ* temperature. In total, the bacterial communities of Axial and Magic Mountain were more similar to each other than to the communities of Kama’ehuakanaloa (Figure 3). Three of the four Axial MGCs incubated at Marker N4 clustered together with AxMGC-47 separated from AxMGC-38 and AxMGC-36. The other marker N4 Axial MGC was incubated for two days, AxMGC-01, and clustered with Axial MGCs from Marker 33 and one of the MGCs from Magic Mountain.

**Figure 3.**
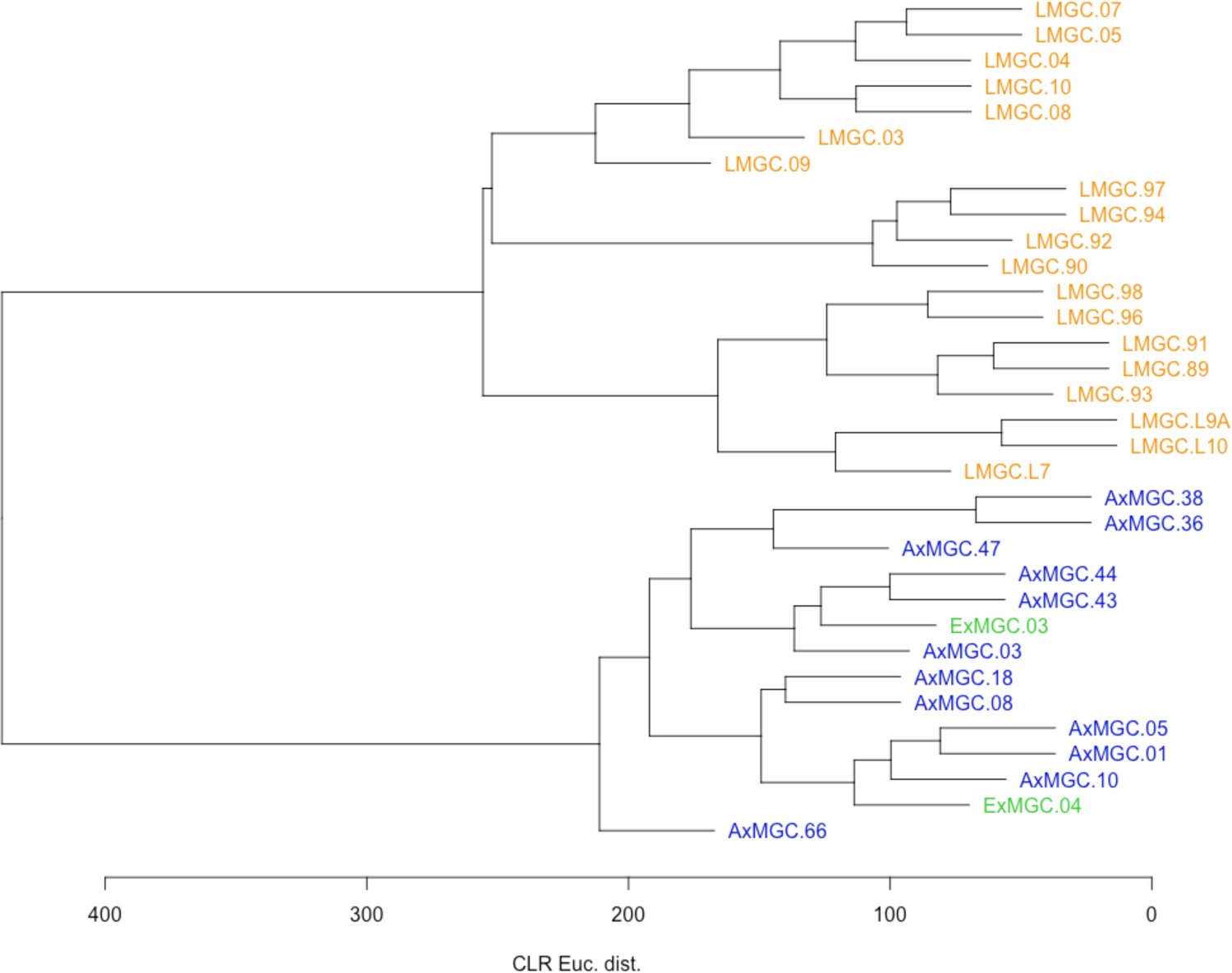
Hierarchical clustering of MGCs communities colored by *in situ* incubation location.

The four Marker 57 (Pohaku) MGCs formed a distinct cluster from the other MGCs at Kama’ehuakanaloa (e.g., LMGC-90, −92, −94, −97). Interestingly, two of the Marker 38 MGCs, LMGC-05 and LMGC-07, clustered separately from the other Marker 38 MGC, LMGC-96. This same pattern was observed for MGCs incubated at Marker 36. The three MGCs from Hiolo North (Markers 36 and 39) clustered together for *in situ* incubations in 2009 but not in 2004. The three samples from Kama’ehuakanaloa incubated shortly after the eruptive event in 1996 formed a unique cluster even though LMGC-L7 was incubated at a cooler temperature than LMGC-L10 and LMGC-L9A.

Duration and temperature were not able to be tested by permanova analysis because the assumption of homogenous within-group dispersions was not met. However, this criterion was met for seamount location and unsurprisingly is significant (p=0.001). Further analysis was completed on Axial separate from Magic Mountain and Kama’ehuakanaloa. Temperature, duration, marker, and year were not significantly correlated with the MGC communities at Axial. However, when preforming permanova on the Kama’ehuakanaloa MGCs to the exclusion of Axial and Magic Mountain, duration (p=0.011) and year (p=0.044) were significant factors in community composition. Magic Mountain microbial communities are represented by only two MGCs, therefore permanova analysis could not be performed.

To further investigate geochemical parameters as drivers of the MGC community structure, permanova analysis was performed (Supplemental Table 2). Both duration and marker were unable to be tested since the within-group dispersions were significant. By permanova, hydrogen sulfide (p=0.385), pH (p=0.069) and manganese (p=0.174) were all non-significant factors in driving community composition. Whereas seamount (p=0.001), year (p=0.001), temperature (p=0.01), and dissolved iron (p=0.001) were significant (Supplemental Table 2). Therefore, redundancy analysis was completed on the top ten taxa by class and significant chemistry parameters. From this, 57.9% of the variance was captured by RDA1 and RDA2 (Figure 4). Dissolved iron and year were the most important factors in RDA1 and abundance of Zetaproteobacteria correlated with the dissolved iron concentration. By RDA analysis, H_2_S and Mn had inverse impacts on community structure than Fe and pH (Figure 4). All locations with iron-dominated fluids from Kama’ehuakanaloa grouped with the Zetaproteobacteria, whereas the hot temperature MGCs from Kama’ehuakanaloa were ordinated with temperature and Aquificae (Figure 4).

**Figure 4.**
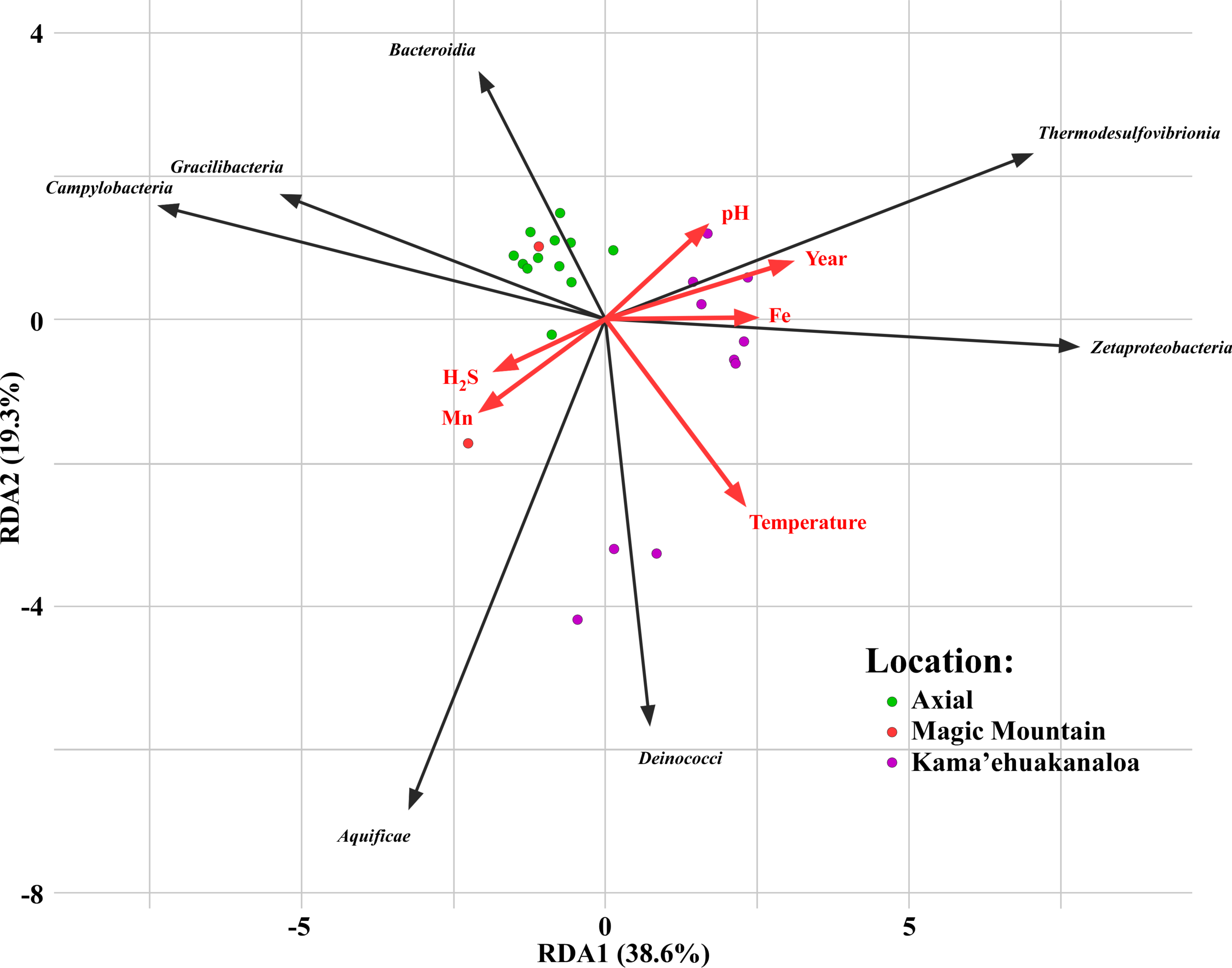
RDA of top 10 most abundant bacterial classes within the MGCs at the three venting locations with selected environmental parameters that were shown to be significant by permanova.

Within the Campylobacteria, the genera *Sulfurovum* and *Sulfurimonas* were present in all MGCs from Axial and Magic Mountain (Figure 5A). Nine of the twelve MGCs from Axial were dominated by *Sulfurovum* and the other three were dominated by *Thioreductor* and *Sulfurimonas*. On average, Campylobacteria accounted for 77.43% ± 16.6% of the ASVs across the MGCs from Axial and Magic Mountain. AxMGC-05 was 95.6% Campylobacteria, which was mostly *Sulfurovum* and *Sulfurimonas*. Only one sample from Axial or Magic Mountain had less than 50% Campylobacteria: AxMGC-66 with 36.62%, which was primarily *Sulfurovum*. That MGC also had the longest *in situ* incubation time. Based on previous research at Axial, dominance of these known sulfur-oxidizers was expected. However, the Axial MGCs show increased variability by taxa and ASVs even at small geographic distances. Zetaproteobacteria composed less than 0.05% of the total reads in the MGCs incubated Axial and Magic Mountain and were completely absent in four Axial MGCs (AxMGC-05, AxMGC-10, AxMGC-43, and AxMGC-44) and one Magic Mountain MGC (ExMGC-04). Those Zetaproteobacteria that were identified were predominately zOTU-01 and zOTU-02.

**Figure 5.**
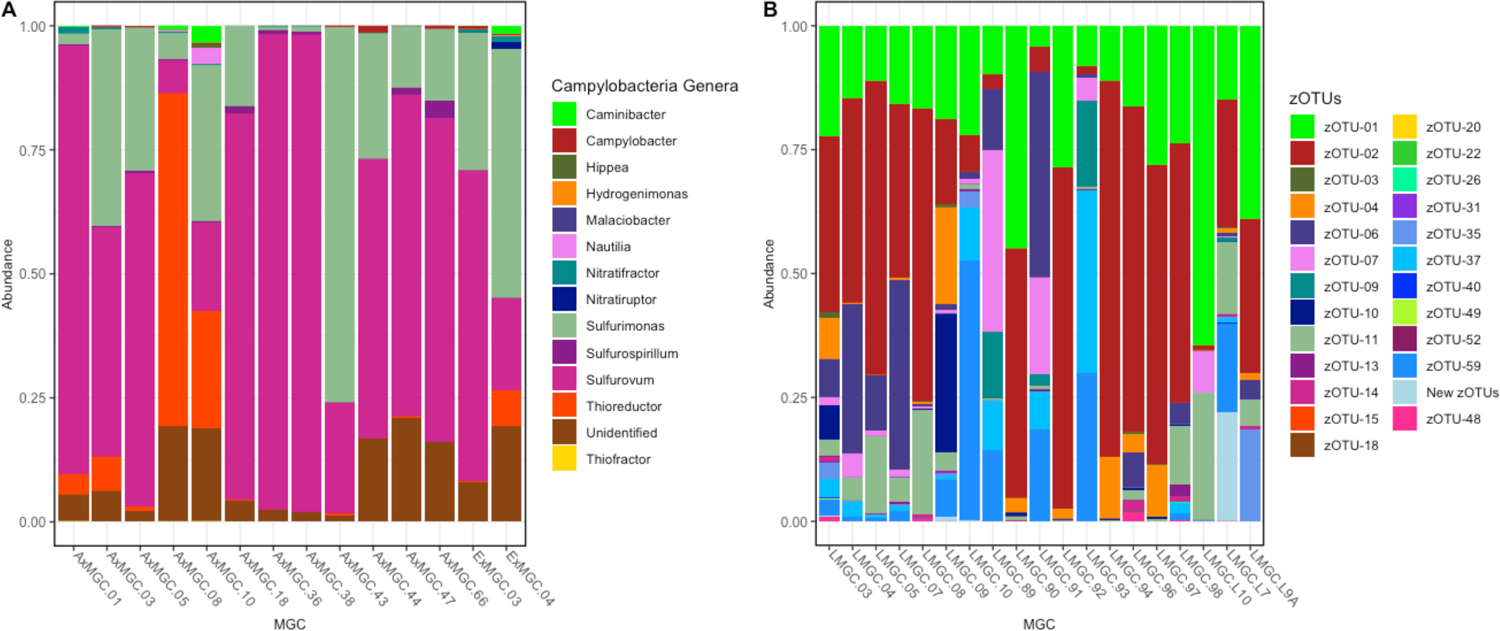
Stacked bar graphs showing (A) abundance of Campylobacteria genera at Axial Seamount and Explorer Ridge and (B) abundance of zOTUs at Kamaʻehuakanaloa Seamount.

In contrast, the MGCs from Kama’ehuakanaloa had a more variable Campylobacteria composition with three MGCs having abundant *Nitratiruptor* and six where *Sulfurimonas* was most dominant. Across all the MGCs from Kama’ehuakanaloa, Campylobacteria accounted for 13.83% of the total community and eight had less than 0.5% Campylobacteria. Zetaproteobacteria ranged in total population of the Kama’ehuakanaloa MGCs from 98.99% in LMGC-90 to only 2.81% in LMGC-L9A. zOTU-01 and zOTU-02 were present in all the Kama’ehuakanaloa MGCs and in eleven of these MGCs, zOTU-01 and zOTU-02 composed 250% of the total Zetaproteobacteria amplicons (Figure 5B). Three MGCs had a high abundance of −New zOTUs”, which represent Zetaproteobacteria sequences that have not been classified into a previously identified zOTU (McAllister et al., 2018). Representatives of zOTUs 3, 9, 11, 14, 18, 23, 36 and 37 have cultured representatives, and only zOTUs 23 and 36 were not identified within the Kama’ehuakanaloa MGCs. Due to the high levels of reduced iron present at Kama’ehuakanaloa, it is unsurprising that Zetaproteobacteria were the most abundant class.

### Fine-scale community analysis

A subset of the most dominant ASVs was further analyzed to better understand persistence among incubation timeframes. From the heatmap shown in Figure 6, there is a clear distinction between the Kama’ehuakanaloa MGCs and the MGCs from Axial and Magic Mountain Furthermore, the Kama’ehuakanaloa MGCs incubated after the eruptive event are also distinct from all other MGCs. Abundant ASVs in the Magic Mountain and Axial Seamount MGCs had very low abundances at Kama’ehuakanaloa and conversely, the abundant ASVs from Kama’ehuakanaloa had low abundances in the Magic Mountain and Axial MGCs. Only ASV-6, a *Sulfurimonas* spp., showed similar abundances across all MGCs locations. ASV-56, also a *Sulfurimonas* spp., also showed high abundance at Kama’ehuakanaloa but was largely absent from Axial and Magic Mountain MGCs. zOTUs 1, 2, 6, 7, 11, 37 and 59 were represented in the most abundant ASVs (Supplemental Table 3). Cultured representatives of the Zetaproteobacteria were found in the MGC community, but none of them were identified in the topmost abundant ASVs.

**Figure 6.**
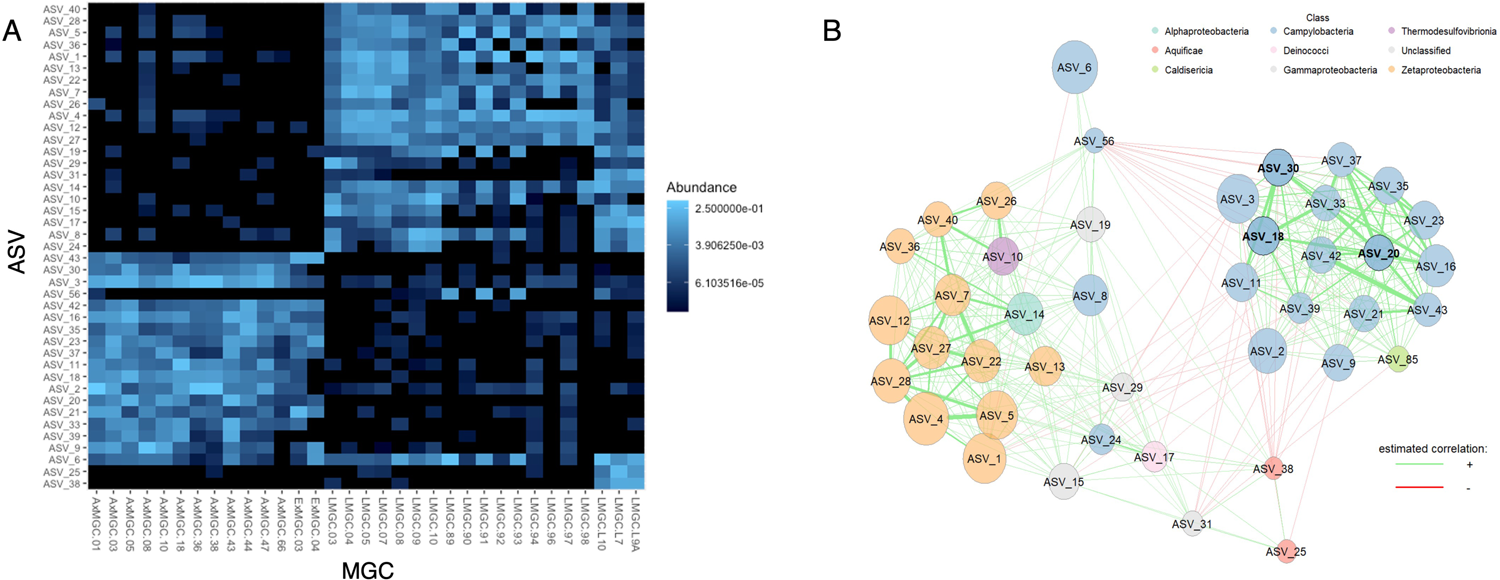
(A) Heatmap displaying abundance of the top 0.5% ASV within the MGCs from Axial Seamount, Explorer Ridge, and Kamaʻehuakanaloa Seamount and (B) their interaction network. Color of nodes corresponds to bacterial class; edge colors correspond to positive (green) or negative (red) estimated correlation. Bold text indicates that node acts as a hub.

An association network was constructed to visualize interactions among the most abundant ASVs. Two discrete clusters were formed within the network joined by strong mutual positive correlations and separated by weakly negative correlations between multiple ASVs within each cluster (Figure 6B). One cluster was composed almost entirely of members of class Campylobacteria that were highly present in both Axial and Magic Mountain communities while having low presence in Kama’ehuakanaloa MGC communities. The other cluster was largely composed of members of class Zetaproteobacteria with strong within-class positive correlations, but also included Campylobacteria, Aquificae, and Gammaproteobacteria, all of which were highly represented at Kama’ehuakanaloa. Regardless of cluster, the strongest correlations were positive, but several ASVs in the Kama’ehuakanaloa-dominated cluster correlated negatively to multiple ASVs in the Axial-Magic Mountain cluster. Interestingly, ASV-56, a *Sulfurimonas* spp. in the Kama’ehuakanaloa cluster (and found only in Kama’ehuakanaloa MGCs), exhibited the most abundant negative correlations, all to the Campylobacteria present in the Axial-Magic Mountain Campylobacteria cluster.

Of the Zetaproteobacteria present in the network analysis, zOTU-01 is represented by two distinct ASVs (ASV-5 and ASV-12) and zOTU-02 is represented by five different ASVs (ASV-1, ASV-4, ASV-22, ASV-27, and ASV-28). These ASVs showed a strong positive relationship to each other and to ASV-14, an Alphaproteobacteria. The remainder of the zOTUs in this analysis are represented by a single ASV and were identified as zOTU-06, zOTU-07, zOTU-11, zOTU-37, and zOTU-59. Two cultured Zetaproteobacteria are present within this analysis, zOTU-11 and zOTU-37, which are represented by ASV-14 and ASV-40, respectively. Interestingly, the type strain of Zetaproteobacteria, zOTU-11, does not form strong relationships with any other ASVs. However, zOTU-37 forms a strong relationship with other Zetaproteobacteria and a Thermodesulfovibrio.

ASV-6, the only highly abundant Campylobacteria found in all three vent locations (Figure 6A) clustered with Kama’ehuakanaloa Seamount but failed to show any strong correlations with other taxa (Figure 6B). Three network hubs, representative of the most influential taxa in the network, were identified. All were Campylobacteria that were only found in the Axial-Magic Mountain cluster. Unsurprisingly, there were no positive correlations between the Zetaproteobacteria of the Kama’ehuakanaloa Seamount cluster and any of the Axial-Magic Mountain cluster ASVs, even though some Zetaproteobacteria were identified as minor community members of the MGCs from Axial and Magic Mountain.

## Discussion

MGCs were placed within hydrothermal vent effluent to better understand the dynamics of microbial colonization at three Pacific Ocean hydrothermal vent sites. These incubations showed distinct microbial communities based on vent location, which relate to the predominant chemical composition of the discharging hydrothermal fluid. It is unclear if the nascent communities are developed from the dominant members of the existing microbial communities at these vents or if they are stochastically colonized. Alternatively, these nascent communities within the MGCs could represent ASVs that are adapted to growing on new surfaces. Additionally, shortly after the Kama’ehuakanaloa eruptive event, the colonizing community had a greater abundance of Campylobacteria as compared to the incubations at cooler temperatures, which were dominated by Zetaproteobacteria. This shows that both temperature and location of the vents has an influence on the microbial community composition. When including the metadata collected alongside the MGCs, vent location, year, temperature, and dissolved iron were significant by permanova analysis. Changes in community composition observed between sampling years could be due to the ephemeral nature of the discharging water (e.g., temperature, composition, flow rate), or due to different *in situ* incubation durations. It is unsurprising that dissolved iron was a driver of community structure since the zOTUs identified within the MGCs are hypothesized to be obligate iron-oxidizers and all tested sites had measurable amounts of dissolved iron.

### Overall Diversity and Community Differences Across Locations

Overall, the MGCs showed no pattern of diversity as related to duration of incubation or incubation temperatures. The year of MGC incubation was a significant factor for community composition in the MGCs, which is unsurprising due to the dynamic nature of these environments. However, even with this variation, the same ASVs are present within the MGCs per location, although their abundance varies between years of incubation and they have distinct ASVs as dominant community members. A previous study at the East Pacific Rise 13° N showed more phylotypes in short-term colonization experiments as compared to the longer-term incubations at the same site (Alain *et al*. 2004). In comparison to other deep-sea studies on microbial communities, we do not see patterns of succession as observed at whale and wood fall sites (Goffredi and Orphan 2010; Kalenitchenko *et al*. 2016). This could be due to the relatively short incubation times or the constant influx of reduced substrates in hydrothermal fluids.

Other deep-sea colonization experiments performed on rock-chips have noted evidence of a changing community over time and microbially-driven chemical weathering (Orcutt *et al*. 2011a; Gulmann *et al*. 2015). Previous studies have shown a higher diversity in mature iron-dominated microbial mats than those that are sulfur-dominated in the Mariana Arc and back-arc (Hager *et al*. 2017). However, this pattern of higher diversity in high dissolved iron sites was not observed in the MGC microbial communities. Microbial mat surfaces, which are hypothesized to contain the active fraction in mature mats, show a wider range in richness as measured by Chao1, from 2,140 to 31,580 (Chan *et al*. 2016; Duchinski *et al*. 2019). This indicates that the MGCs are lower in taxonomic diversity than that of established microbial mats. With the decreased microbial diversity of nascent mat communities, as represented by the MGCs, there is likely to be less functional diversity than that found in established communities as well (Wagg *et al*. 2021).

Gammaproteobacteria that were identified from Kama’ehuakanaloa MGCs have also been found in established microbial mats, though the lack of these taxa within the Axial and Magic Mountain MGCs is in contrast to another previous study (Opatkiewicz, Butterfield and Baross 2009; Rassa *et al*. 2009). The nascent community composition from Axial Seamount and Magic Mountain was similar to active hydrothermal chimneys from Manus Basin (Meier *et al*. 2017, 2019) and vent fluids from Brother’s Volcano and the East Pacific Rise (Zhou *et al*. 2022).

### Zetaproteobacteria Diversity

Both Campylobacterota and Zetaproteobacteria are chemolithoautotrophs and can support a diversity of microorganisms and macrofauna. Genomic and proteomic analysis of Zetaproteobacteria show autotrophy is supported by the Calvin-Benson-Bassham cycle (Singer *et al*. 2011; Barco *et al*. 2015). Conversely, Campylobacterota and Aquificota primarily use the reverse TCA cycle for carbon fixation (Hügler and Sievert 2011). Analysis of carbon fixation genes from established microbial mats show no relationship between Zetaproteobacteria and genes for the rTCA cycle (Jesser *et al*. 2015). Zetaproteobacteria have been shown to produce twisted iron-oxyhydroxide biominerals, which contribute to the mat architecture (Chan, Emerson and Luther 2016). Within the MGCs, zOTU-01 zOTU-02 were the most dominant, which is expected since they have been found in high abundance within established mats at iron-dominated hydrothermal vent communities (McAllister *et al*. 2019). The metabolism of these dominant zOTUs has been inferred from single cell-genomes and metagenome-assembled genomes (Field *et al*. 2015; Fullerton *et al*. 2017), and both show the capacity for living in aerobic conditions and fixing carbon. These zOTUs also contain a fused cytochrome-porin that is responsible for the oxidation of dissolved Fe^2+^ (Keffer *et al*. 2021), which is conserved across the genus *Mariprofundus* (Zhong *et al*. 2022). It is unlikely that the Zetaproteobacteria are growing via hydrogen oxidation since only zOTU-09 is known with this metabolism (Mori *et al*. 2017) and was present as minor community members within six of the 19 MGCs incubated at Kama’ehuakanaloa.

### Campylobacteria Composition

Diverse genera within the phylum Campylobacterota have been shown to inhabit hydrothermal vent plumes, sulfides, and the vent subsurface (Sylvan *et al*. 2012; Dick *et al*. 2013; Meyer *et al*. 2013b). Overall, Campylobacterota were nearly universal within this study, but genus varied between locations. Even in the high temperature Kama’ehuakanaloa Seamount MGCs, the Campylobacterota ASVs were not like those found at Axial or Magic Mountain, as was the case for ASV-56 (*Sulfurimonas* spp.), which was found only at Kama’ehuakanaloa. It was not surprising that *Sulfurimonas* and *Sulfurovum* were present as dominant representatives of Campylobacterota at Kama’ehuakanaloa and were also present in Axial and Magic Mountain MGC’s; due to their diverse metabolisms, they can live in various conditions by using different energy sources (Nakagawa *et al*. 2005; Campbell *et al*. 2006). These groups of bacteria are not only able to use sulfur and iron, but also have a pathway for NO ^-^ reduction (Han and Perner 2015; Pérez-Rodríguez *et al*. 2017). Where there was a dominance of Campylobacteria, the Zetaproteobacteria were minor community members. The high abundance of Campylobacteria as well as Aquificae--a similar phylum of thermophiles that can tolerate higher temperatures and use hydrogen as the electron donor (Reysenbach 2001)--at Kama’ehuakanaloa after the eruption is representative of how the geochemistry of the active vent shifted the microbial community. This is in contrast to other studies from 1998-99 where the diversity of Campylobacteria decreased (Huber, Butterfield and Baross 2003).

### Community interactions

Overall, our results indicate that the concentration of dissolved iron from the vent fluids favor the growth of Zetaproteobacteria as the dominant primary producer. The influence of pH, Mn and H_2_S were not significant in these communities. This is consistent with the RDA analysis and the community interaction, which indicate that the bacterial community is significantly affected by the abundance of Zetaproteobacteria and with their role as ecosystem engineers (Chan *et al*. 2011; Hager *et al*. 2017; Beam *et al*. 2018b).

Although the two classes, Zetaproteobacteria and Campylobacteria, are foundational members of these nascent communities, only the Campylobacteria formed hubs in network analysis, which indicate keystone taxa (Peschel *et al*. 2021). Two ASVs from class Aquificae formed positive associations with the Zetaproteobacteria in the Kama’ehuakanaloa cluster, despite their lack of cooccurrence in time, while also forming negative associations with the Campylobacteria from the Axial-Magic Mountain cluster despite the earlier noted tendency for Aquificota and Campylobacteria to cooccur. These seemingly contradictory associations further support the significance of geographic location as a driver of community structure. Within the Zetaproteobacteria, the most influential community members were identified as belonging to zOTU-06 (ASV_7) and zOTU-02 (ASV_27, ASV_22 and ASV_28) (Supplemental Table 2). However, they did not meet the cutoff requirement for hub classification and so are not considered as keystone taxa, which is in contrast to other iron-rich marine environments (Beam *et al*. 2018a). Additionally, the Gammaproteobacteria are found throughout the nascent communities at Axial, Magic Mountain and Kama’ehuakanaloa. Only a few Gammaproteobacteria ASVs formed positive relationships with foundational members of those communities, and only with the Zetaproteobacteria.

## Conclusion

This study builds on the knowledge of community diversity and composition of lithotrophic bacteria, providing new data from three Pacific Ocean hydrothermal vent locations. Our results indicate that diverse microbial communities form quickly within hydrothermal vent effluent and are dominated by chemolithoautotrophs. SSU rRNA amplicon sequencing from 33 MGCs showed relatively low alpha diversity overall. There was also no difference in diversity across incubation periods and each site showed distinct microbial community composition. This supports the influence of biogeography as well as the impact of geochemistry on microbial diversity across these vent locations. Although Shannon and Simpson’s diversity indices did not show correlation with age of MGC, future comparisons should be performed between the mature and nascent microbial mat to further understand the drivers of diversity.

Also of note was the reduction in the abundance of iron-oxidizing Zetaproteobacteria after the Kama’ehuakanaloa eruption, which instead displayed a higher abundance of Campylobacteria. Observing such shifts is important to understanding how the biogeochemistry of the surrounding area affects autotrophic microbial communities. Additionally, further research should focus on determining if the community members present within the MGCs are also present in the vent fluids or if they are members of the adjacent microbial mats.

Future studies should include longer incubation periods to further determine whether patterns of succession are different for these habitats as opposed to whale and wood falls. In addition, a simultaneous study of vent fluids, established microbial mats, and MGC composition should be conducted to understand whether the community composition is the same across these substrates under the same incubation conditions.

## Author Contribution

HF and CLM designed the work. CLM was responsible for sample collection. HF conducted most of the data analysis. LS is responsible for creation of network diagrams. HF, LS, AE, and CLM wrote the manuscript and approved the final version to be published. Corresponding geochemical assays for Axial and Magic Mountain were designed and conducted by DB and by CGW for Kama’ehuakanaloa Seamount. HF, LS, AE, DB, CGW and CLM were responsible for all aspects of this work and approved the published version.

## Funding

This project was funded in part by National Science Foundation awards MCB-0348734 (to CLM) and the College of Charleston Department of Biology Research and Development fund (to HF).

## Supporting information

similar patterns to overall observed ASVs (Supplemental Table 1). Both alpha diversity estimators

permanova analysis was performed (Supplemental Table 2). Both duration and marker were unable

abundant ASVs (Supplemental Table 3). Cultured representatives of the Zetaproteobacteria were

## Acknowledgments

We wholeheartedly thank the operation teams for Pisces V and Jason II, and the captains and crew of the R/Vs Kaimikai-o-Kanaloa, Melville, Kilo Moana, and Thomas G. Thompson for their assistance with sample collection during oceanographic research cruises in 1998, 2000, 2002, 2004 and 2009.

## Notes

### Competing Interest Statement

The authors have declared no competing interest.

