## Supplementary material for "Seafloor Incubation Experiments at Deep-Sea Hydrothermal Vents Reveal Distinct Biogeographic Signatures of Autotrophic Communities": similar patterns to overall observed ASVs (Supplemental Table 1). Both alpha diversity estimators

**Supplemental Table 1:** Diversity indices of MGC from three Seamounts.

| MGC | Duration | Observed | Chao1 | ACE | Shannon | Simpson | InvSimpson | Fisher |
|---|---|---|---|---|---|---|---|---|
| AxMGC.01 | 2 | 117 | 117.00 | 117.23 | 2.20 | 0.66 | 2.92 | 13.47 |
| AxMGC.03 | 4 | 1278 | 1300.79 | 1302.21 | 4.06 | 0.95 | 21.04 | 165.92 |
| AxMGC.05 | 16 | 102 | 102.00 | 102.00 | 3.02 | 0.91 | 11.43 | 12.02 |
| AxMGC.08 | 2 | 840 | 863.82 | 863.36 | 3.18 | 0.81 | 5.19 | 110.08 |
| AxMGC.10 | 14 | 186 | 186.00 | 186.00 | 3.81 | 0.95 | 20.57 | 22.46 |
| AxMGC.18 | 12 | 861 | 883.99 | 883.85 | 3.15 | 0.86 | 7.38 | 108.08 |
| AxMGC.36 | 5 | 983 | 1005.17 | 1008.19 | 2.26 | 0.71 | 3.47 | 118.21 |
| AxMGC.38 | 5 | 626 | 643.15 | 642.75 | 2.00 | 0.68 | 3.16 | 75.86 |
| AxMGC.43 | 5 | 449 | 473.89 | 470.37 | 3.00 | 0.91 | 10.72 | 55.81 |
| AxMGC.44 | 7 | 617 | 636.13 | 636.08 | 3.75 | 0.95 | 18.23 | 78.40 |
| AxMGC.47 | 7 | 843 | 862.57 | 858.04 | 4.48 | 0.96 | 25.60 | 122.65 |
| AxMGC.66 | 19 | 1144 | 1158.59 | 1159.31 | 4.81 | 0.97 | 33.32 | 165.13 |
| ExMGC.03 | 4 | 544 | 554.32 | 555.09 | 3.81 | 0.94 | 15.44 | 70.78 |
| ExMGC.04 | 4 | 140 | 140.00 | 140.00 | 3.50 | 0.95 | 18.84 | 15.95 |
| LMGC.03 | 4 | 1791 | 1813.36 | 1812.58 | 4.12 | 0.94 | 15.96 | 266.72 |
| LMGC.04 | 4 | 1012 | 1044.27 | 1041.75 | 3.60 | 0.93 | 14.49 | 144.65 |
| LMGC.05 | 4 | 947 | 982.04 | 975.67 | 3.38 | 0.92 | 12.68 | 126.86 |
| LMGC.07 | 4 | 1534 | 1557.26 | 1560.78 | 3.83 | 0.93 | 13.75 | 203.28 |
| LMGC.08 | 4 | 1022 | 1065.50 | 1058.47 | 3.67 | 0.94 | 16.64 | 136.36 |
| LMGC.09 | 4 | 1728 | 1767.46 | 1762.91 | 4.85 | 0.95 | 19.84 | 232.29 |
| LMGC.10 | 4 | 996 | 1024.73 | 1025.06 | 3.28 | 0.89 | 9.44 | 132.72 |
| LMGC.89 | 5 | 202 | 202.11 | 202.72 | 2.73 | 0.84 | 6.30 | 21.47 |
| LMGC.90 | 8 | 1382 | 1421.08 | 1409.04 | 2.95 | 0.82 | 5.47 | 178.78 |
| LMGC.91 | 5 | 107 | 107.50 | 107.45 | 3.17 | 0.93 | 13.43 | 13.03 |
| LMGC.92 | 9 | 1074 | 1121.67 | 1105.38 | 2.35 | 0.77 | 4.39 | 134.73 |
| LMGC.93 | 8 | 242 | 242.14 | 242.45 | 3.65 | 0.95 | 21.16 | 28.55 |
| LMGC.94 | 8 | 931 | 954.46 | 950.85 | 2.73 | 0.82 | 5.53 | 115.18 |
| LMGC.96 | 8 | 129 | 129.25 | 129.55 | 3.52 | 0.94 | 16.47 | 17.67 |
| LMGC.97 | 9 | 756 | 785.06 | 777.61 | 2.79 | 0.84 | 6.11 | 98.52 |
| LMGC.98 | 9 | 281 | 281.00 | 281.00 | 3.86 | 0.96 | 25.90 | 33.30 |
| LMGC.L10 | 4 | 339 | 353.50 | 351.50 | 2.85 | 0.87 | 7.97 | 38.41 |
| LMGC.L7 | 4 | 796 | 819.25 | 822.71 | 3.58 | 0.94 | 16.93 | 108.79 |
| LMGC.L9A | 4 | 207 | 226.89 | 226.69 | 2.91 | 0.90 | 9.66 | 24.22 |
