## Supplementary material for "Seafloor Incubation Experiments at Deep-Sea Hydrothermal Vents Reveal Distinct Biogeographic Signatures of Autotrophic Communities": permanova analysis was performed (Supplemental Table 2). Both duration and marker were unable

**Supplemental Table 2**: Summary of the PERMANOVA analyses based on euclidean distance of ASVs for bacterial communities for the MGCs with associated geochemical measurements

| **Factor** | **p-value for Group Dispersions** | **p-value for Permutational ANOVA** |
|---|---|---|
| Duration | 0.003** | NA |
| Seamount | 0.175 | 0.001*** |
| Year | 0.353 | 0.001*** |
| Marker | 0.047* | NA |
| Temperature | 0.241 | 0.01** |
| pH | 0.157 | 0.069 |
| $H_2S$ | 0.064 | 0.385 |
| dFe | 0.157 | 0.001*** |
| Mn | 0.17 | 0.174 |
