## Supplementary material for "Seafloor Incubation Experiments at Deep-Sea Hydrothermal Vents Reveal Distinct Biogeographic Signatures of Autotrophic Communities": abundant ASVs (Supplemental Table 3). Cultured representatives of the Zetaproteobacteria were

**Supplemental Table 3:** Taxomonic identification of most abundant taxa

| ASV | domain | phylum | class | order | family | genus | zOTU |
|---|---|---|---|---|---|---|---|
| ASV_1 | Bacteria | Proteobacteria | Zetaproteobacteria | Mariprofundales | Mariprofundaceae | Mariprofundus | zOTU-02 |
| ASV_10 | Bacteria | Nitrospirota | Thermodesulfovibrionia | Thermodesulfovibrionales | Thermodesulfovibrionaceae | Thermodesulfovibrio | |
| ASV_11 | Bacteria | Campilobacterota | Campylobacteria | Campylobacterales | Sulfurovaceae | Sulfurovum | |
| ASV_12 | Bacteria | Proteobacteria | Zetaproteobacteria | Mariprofundales | Mariprofundaceae | Mariprofundus | zOTU-01 |
| ASV_13 | Bacteria | Proteobacteria | Zetaproteobacteria | Mariprofundales | Mariprofundaceae | Mariprofundus | zOTU-11 |
| ASV_14 | Bacteria | Proteobacteria | Alphaproteobacteria | Rhodobacterales | Rhodobacteraceae | Roseobacter clade Marinomonas lineage | |
| ASV_15 | Bacteria | NA | NA | NA | NA | NA | |
| ASV_16 | Bacteria | Campilobacterota | Campylobacteria | Campylobacterales | Sulfurovaceae | Sulfurovum | |
| ASV_17 | Bacteria | Deinococcota | Deinococci | Thermales | Thermaceae | Vulcanithermus | |
| ASV_18 | Bacteria | Campilobacterota | Campylobacteria | Campylobacterales | Sulfurovaceae | Sulfurovum | |
| ASV_19 | Bacteria | Proteobacteria | Gammaproteobacteria | Thiomicrospirales | Thiomicrospiraceae | Galenea | |
| ASV_2 | Bacteria | Campilobacterota | Campylobacteria | Campylobacterales | Sulfurovaceae | Sulfurovum | |
| ASV_20 | Bacteria | Campilobacterota | Campylobacteria | Campylobacterales | Sulfurimonadaceae | Sulfurimonas | |
| ASV_21 | Bacteria | Campilobacterota | Campylobacteria | Campylobacterales | Sulfurovaceae | Sulfurovum | |
| ASV_22 | Bacteria | Proteobacteria | Zetaproteobacteria | Mariprofundales | Mariprofundaceae | Mariprofundus | zOTU-02 |
| ASV_23 | Bacteria | Campilobacterota | Campylobacteria | Campylobacterales | Sulfurimonadaceae | Sulfurimonas | |
| ASV_24 | Bacteria | Campilobacterota | Campylobacteria | Campylobacterales | Nitratiruptoraceae | Nitratiruptor | |
| ASV_25 | Bacteria | Aquificota | Aquificae | Aquificales | Aquificaceae | Hydrogenivirga | |
| ASV_26 | Bacteria | Proteobacteria | Zetaproteobacteria | Mariprofundales | Mariprofundaceae | Mariprofundus | zOTU-59 |
| ASV_27 | Bacteria | Proteobacteria | Zetaproteobacteria | Mariprofundales | Mariprofundaceae | Mariprofundus | zOTU-02 |
| ASV_28 | Bacteria | Proteobacteria | Zetaproteobacteria | Mariprofundales | Mariprofundaceae | Mariprofundus | zOTU-02 |
| ASV_29 | Bacteria | Proteobacteria | Gammaproteobacteria | Pseudomonadales | Pseudomonadaceae | Pseudomonas | |
| ASV_3 | Bacteria | Campilobacterota | Campylobacteria | Campylobacterales | Sulfurovaceae | Sulfurovum | |
| ASV_30 | Bacteria | Campilobacterota | Campylobacteria | Campylobacterales | Sulfurovaceae | Sulfurovum | |
| ASV_31 | Bacteria | Proteobacteria | Gammaproteobacteria | Nitrococcales | Halorhodospiraceae | Sulfurivirga | |
| ASV_33 | Bacteria | Campilobacterota | Campylobacteria | Campylobacterales | Sulfurimonadaceae | Sulfurimonas | |
| ASV_35 | Bacteria | Campilobacterota | Campylobacteria | Campylobacterales | Sulfurovaceae | Sulfurovum | |
| ASV_36 | Bacteria | Proteobacteria | Zetaproteobacteria | Mariprofundales | Mariprofundaceae | Mariprofundus | zOTU-07 |
| ASV_37 | Bacteria | Campilobacterota | Campylobacteria | Campylobacterales | Sulfurimonadaceae | Sulfurimonas | |
| ASV_38 | Bacteria | Aquificota | Aquificae | Aquificales | Aquificaceae | Aquifex | |
| ASV_39 | Bacteria | Campilobacterota | Campylobacteria | Campylobacterales | Sulfurimonadaceae | Sulfurimonas | |
| ASV_4 | Bacteria | Proteobacteria | Zetaproteobacteria | Mariprofundales | Mariprofundaceae | Mariprofundus | zOTU-02 |
| ASV_40 | Bacteria | Proteobacteria | Zetaproteobacteria | Mariprofundales | Mariprofundaceae | Mariprofundus | zOTU-37 |
| ASV_42 | Bacteria | Campilobacterota | Campylobacteria | Campylobacterales | Sulfurovaceae | Sulfurovum | |
| ASV_43 | Bacteria | Campilobacterota | Campylobacteria | Campylobacterales | Sulfurovaceae | Sulfurovum | |
| ASV_5 | Bacteria | Proteobacteria | Zetaproteobacteria | Mariprofundales | Mariprofundaceae | Mariprofundus | zOTU-01 |
| ASV_56 | Bacteria | Campilobacterota | Campylobacteria | Campylobacterales | Sulfurimonadaceae | Sulfurimonas | |
| ASV_6 | Bacteria | Campilobacterota | Campylobacteria | Campylobacterales | Sulfurimonadaceae | Sulfurimonas | |

| | | | | | | | |
|---|---|---|---|---|---|---|---|
| ASV_7 | Bacteria | Proteobacteria | Zetaproteobacteria | Mariprofundales | Mariprofundaceae | Mariprofundus | zOTU-06 |
| ASV_8 | Bacteria | Campilobacterota | Campylobacteria | Campylobacterales | Nitratiruptoraceae | Nitratiruptor | |
| ASV_9 | Bacteria | Campilobacterota | Campylobacteria | Campylobacterales | Thioreductoraceae | Thioreductor | |
